# Direct Intercellular Vesicle Exchange between Adjacent Cells

**DOI:** 10.1101/2024.12.11.626728

**Authors:** Tomohiro Minakawa, Tatsuya Katsuno, Keiko Okamoto-Furuta, Ryoko Ando, Fumiyoshi Ishidate, Takahiro K. Fujiwara, Dooseon Cho, Sho Takehana, Yoshikatsu Sato, Yasuhiko Tabata, Atsushi Miyawaki, Jun K. Yamashita

## Abstract

Intercellular communication plays a central role in the development and integrity of multicellular organisms. Vesicle transfer, especially through extracellular space, has recently been highlighted as a critical intercellular communication modality, carrying nucleic acids, proteins, and others to distant cells. Previously, we demonstrated that extracellular vesicles induce “phenotypic synchronization of cells (PSyC)” during stem cell differentiation. While examining the mechanism underlying PSyC, we discovered a novel form of cellular communication mediated by direct intercellular vesicle exchange (DIVE) between adjacent cells across the plasma membrane (PM). By achieving cell-wide and high-spatiotemporal resolution imaging of vesicles labeled with fusion proteins of CD63 or CD81 to StayGold, a photostable fluorescent marker, we observed small vesicles (50-500 nm in diameter) directly transferred between adjacent cells. These vesicles moved at approximately 1 µm/s and crossed PM in approximately 10-20 seconds. Furthermore, multiple vesicles traversed nearly identical sites of PM, suggesting the presence of specific routes or structures, potentially including a pore, mediating the vesicle transfer. Three-dimensional electron microscopy provided supportive observations for traversing vesicles with single membrane. These vesicles, named InterCellular Vesicles (InterCVs), were observed to colocalize with nucleic acids, including mRNA, microRNA, and DNA, suggesting the exchange of nucleic acid-mediated information, potentially inducing PSyC, between adjacent cells. Our discovery, DIVE, reveals a previously unknown modality of cell-cell communication, with the potential to reshape our understanding of cellular biology.

## Introduction

Intercellular communication plays a central role in the development and integrity of multicellular organisms, including tissue and organ formation. While there are various mechanisms of intercellular communication, molecular interactions between ligands and their receptors, as well as membrane proteins such as adhesion molecules, channels, and transporters have historically dominated biological research, resulting in a wealth of accumulated knowledge [1]. More recently, it has been discovered that vesicles with diameters ranging from 40 to 1,000 nm, known as extracellular vesicles (EVs), are secreted by cells and can transport nucleic acids, proteins, lipids, and even organelles such as mitochondria to other cells [2, 3]. These findings suggest that EVs convey complex and multifaceted information to recipient cells in the distance, extending beyond simple molecular-level signal transduction [4]. In contrast, relatively little is known about direct communication between adjacent cells. Apart from classical intercellular adhesion structures, such as tight junctions, adherens junctions, and gap junctions [5], many aspects of direct cell-cell communication remain unexplored. Other reports primarily focus on specialized structures such as cellular protrusions. The intercellular bridge (IB), which connects daughter cells at the end of cell division, has long been observed [6]. Recently, its component molecules has been elucidated [7], and it has also been revealed that cytoplasmic material can pass through the IB [8]. Recent discoveries also point to the involvement of tunneling nanotubes (TNTs), which are thin, elongated tubes extending from the cell membrane that connect cells over short distances, enabling the exchange of substances including signaling molecules [9]. Similarly, long protrusions known as cytonemes are observed in cells such as Drosophila wing imaginal disc cells, where they are thought to be involved in morphogen transport [7]. What these discoveries have in common is the necessity of having recognizable structures, even in static observations such as fixed samples. For modalities of intercellular communication that do not exhibit such clear, defined structures, the limitations in capturing dynamic structural changes and real-time behavior of cellular components have been leaving such possibilities largely unexplored.

Previously, we discovered that embryonic stem cells (ESCs) with constitutively active protein kinase A (PKA) differentiated faster than wild-type ESCs [10]. When ESC differentiation was induced under the co-culture of PKA-activated ESCs and wild-type ESCs, which differentiate more slowly, the slower-differentiating cells synchronized their differentiation stage with the faster-differentiating ESCs through accelerating the differentiation speed. Furthermore, we found that EVs were involved in this phenomenon. This synchronization of cells through EVs was named Phenotypic Synchronization of Cells (PSyC) [11]. PSyC was strongly observed during co-culture, but its effect drastically decreased when the cells were cultured separately with a polycarbonate membrane. Based on these results, we focused on the transfer of information, specifically vesicles, between adjacent cells. We attempted real-time monitoring of vesicle movement between adjacent cells. It is challenging to observe small vesicles, such as EVs or other vesicles, by live imaging, and the detailed dynamics of vesicles within and between cells remain poorly understood. Advances in live imaging with higher spatiotemporal resolution technologies have been expected to elucidate the dynamics of small vesicles.

In this study, to observe vesicle dynamics, we introduced fluorescent proteins fused with CD81 or CD63, which are known EV markers, into the cells, and used super-resolution microscopy techniques. By using StayGold (SG), a fluorescent protein that exhibits high brightness and exceptional photostability [12], we were able to monitor vesicles with high spatiotemporal resolution with a considerable extension of the observation period as well as the rapid acquisition of numerous images spanning a broad field. As a result, we identified vesicles moving rapidly within cells at speeds of 1-2.5 µm/s, with some of these vesicles crossing the plasma membrane (PM) between adjacent cells approximately in 10 seconds. Multiple vesicles repeatedly traveled along similar routes and traversed nearly identical sites on PM within tens-of-seconds intervals. These findings led us to hypothesize a new cellular mechanism that guides vesicles to transfer directly into adjacent cells. We named this Direct Intercellular Vesicle Exchange (DIVE). Three-dimensional electron microscopy studies with FIB-SEM (Focused Ion Beam Scanning Electron Microscopy) provided supportive observations for DIVE phenomenon, such as successive vesicles believed to traverse PM and a structure that could potentially correspond to pores in PM. The vesicles exchanged between adjacent cells were considered to be composed of single membrane. We named the vesicles as InterCellular Vesicles (InterCVs). InterCVs carried at least nucleic acids such as microRNA, mRNA, and DNA. DIVE was observed in epithelial cells, differentiating pluripotent stem cells, and even between heterogeneous species of pluripotent stem cells, suggesting that DIVE is broadly present in a wide range of cell types.

In this way, we have discovered a new modality of intercellular communication. Cells may possess more diverse intercellular communication mechanisms than previously thought, and even without specialized structures like protrusions, they may engage in dense exchange of information directly, supporting multicellular systems. While much remains to be explored regarding the detailed structure, mechanisms, biological and evolutionary significance of DIVE, as well as its potential applications in disease understanding and therapy, the discovery of DIVE should be a groundbreaking finding with the potential to reshape our understanding in biology and medicine.

## Results

### Discovery of Vesicles Moving Between Adjacent Cells Through High Spatiotemporal Resolution Live Imaging

First, to visualize vesicles moving between cells, we constructed expression vectors for fusion proteins of fluorescent proteins (FPs) with CD63 or CD81, which are EV markers and are also expressed on various cellular membranes including intracellular vesicles and PM. When fusion proteins of CD63 or CD81 and FPs (Fig. S1A) were introduced into mouse ESCs, both the cell membrane and vesicles were labeled (Fig. S1B). Vesicles could be detected as signals with a diameter of approximately 100-200 nm (Fig. S1C). To distinguish vesicles originating from adjacent cells, we co-cultured mouse ESCs expressing CD81-EGFP (CD81-EGFP-ESC) and those expressing CD81-tdTomato (CD81-tdTomato-ESC) (Fig. S1A, S1B), and examined whether red vesicles could be detected in green cells or vice versa. In the co-cultured cells, we observed green vesicles looking like they were moving in red cell (Fig. S1D, Supplemental movie S1). However, since this data is in two dimensions (single plane), it is unclear whether these vesicles actually entered red cell and moved the inside of red cell. Then, we performed 3D live imaging with a super-resolution confocal microscopy (LSM880 with Airyscan, Carl Zeiss, Germany) and found that several vesicles with nearly linear alignment appeared to move in regions corresponding to PM (Fig. 1A, B), suggesting the transfer of vesicles between adjacent cells. Nevertheless, the spatiotemporal resolution was still insufficient to clearly capture the movement of vesicles between cells, and further optimization of conditions was deemed necessary. A wide field of view in the XY plane (approximately 100 µm x 100 µm) is necessary to reliably capture relatively rare vesicle movement events. To effectively track fast-moving vesicles without losing their trajectories, it is necessary to cover a Z-axis thickness of more than 2.5 µm with Z-step intervals smaller than 300 nm.

**Fig. 1.**
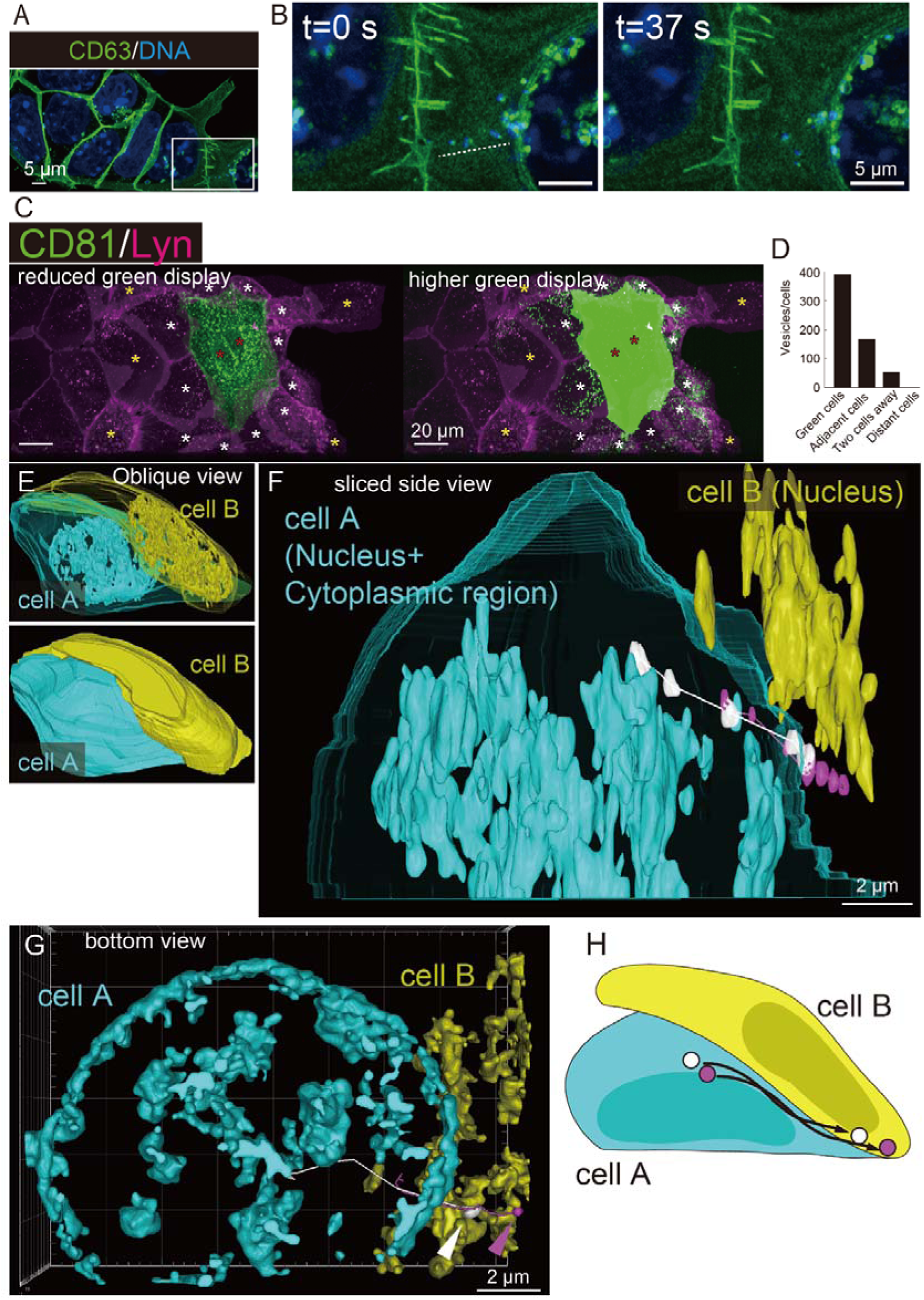
Vesicle transfer between adjacent cells. (**A** and **B**) Vesicles moving in PM region. A confocal live 3D imaging of CD63-EGFP-ESCs. (**A**) Lower magnification view. Green: plasma membrane and vesicles marked with CD63. Blue: DNA marked with DAPI. (**B**) A time series of Box in (A). A linear alignment of vesicles in PM region moving to the left (dotted line). The vesicles co-localized with DNA. Scale bars, 5 μm. (**C** and **D**) Dominant vesicle transfer to adjacent cells (**C**) A 3D image of co-cultured CD81-SG-MDCK (green cells) and Lyn-Azalea-MDCK (red cells). Red asterisks indicate green cells, white asterisks indicate red cells adjacent to green cells, and yellow asterisks indicate red cells two-cells away from green cells, respectively. Scale bars, 20 μm. Higher green intensity image (right panel) clearly visualized green vesicles in red cells. (**D**) A graph showing the number of green vesicles within each cell type. (**E-H**) Vesicle tracking between adjacent cells. CD81-SG-ESCs. Both cell A and cell B were labeled with a new DNA staining dye, PC5 (nuclei) and CD81 (vesicles and PM). Cell A is represented in pseudocolor as blue. Cell B is represented in pseudocolor as yellow. (**E**) Overall positional relationship of cell A and B. Oblique view of 3D structures with cytoplasmic regions shown as transparent (upper panel) and opaque (lower panel). (**F**) Segment sliced side view. The movement and trajectory of two vesicles (white and magenta) from cell A to cell B are overlaid. Scale bar, 2 μm. (**G**) Bottom view of cell A and cell B. Final position of the vesicles in cell B (white and magenta arrowheads, respectively) and trajectories are shown. Scale bar, 2 μm. (**H**) Schematic diagram showing the movement of vesicles (InterCVs: Intercellular vesicles) from cell A to cell B.

Furthermore, capturing a complete 3D image stack within 3 seconds is critical for accurate trajectory analysis. These conditions required the use of strong fluorescence with extremely short exposure times in the millisecond range. In addition, photobleaching observed in conventional FPs after capturing only a few frames have to be overcome. We addressed these issues and achieved the high spatiotemporal resolution necessary for vesicle tracking by combining the highly photostable fluorescent protein StayGold (SG) [12] with a super-resolution SIM microscope (Elyra7 with SIM Apotome, Carl Zeiss, Germany). Conventional FPs with confocal microscopy requires 120 seconds to capture a space of 45×45×2.5 µm in 5 planes (Fig. S1E). In contrast, combining SG with Elyra7 allows capturing a space of 128×128×2.7 µm in 10 planes in just 2.5 seconds (Fig. S1E). Thus, spatiotemporal resolution has been drastically improved.

We introduced CD81-SG into canine kidney epithelial MDCK cells and established CD81-SG-MDCK cells. These cells grow as a single-cell layer without overlapping, making it easier to identify the positions of vesicles (Fig. S1F). To more clearly distinguish the cell boundary with CD81-SG-MDCK cells, we established Lyn-Azalea-MDCK cells, which express a fusion protein consisting of Lyn, a protein localizing to PM, and Azalea, a red FP [13] (Fig. S1F). CD81-SG-MDCK cells and Lyn-Azalea-MDCK cells were seeded at a 1:100 ratio and co-cultured. After 2 days of co-culture, many green vesicles were observed in Lyn-Azalea-MDCK cells adjacent to CD81-SG-MDCK cells. The clear existence of green vesicles in the inside of Lyn-Azalea-MDCK cells was confirmed using a lightsheet microscopy (Fig. S1G, Supplemental movie S2). The number of green vesicles within the surrounding Lyn-Azalea-MDCK cells decreased to less than half for each cell distance away from the CD81-SG-MDCK cells (Fig. 1C, D, Supplemental movie S3), suggesting that there is a vesicle delivery mechanism that dominantly transports vesicles to adjacent cells. We also generated CD81-SG-mESC cells carrying CD81-SG (Fig. S1H) and tried the live imaging of vesicle transfer between adjacent cells. Fig. 1E-H highlighted two cells, cell A (colored light blue) and cell B (colored yellow), both of which nuclei were visualized by a new DNA staining dye, PC5 [14] (Fig. 1E).

Two vesicles were tracked to move from cell A to cell B, apparently passing through the cell A membrane (Fig. 1F, Supplemental movie S4). During this process, the two vesicles looked partially sharing the same pathway in the cell and during crossing PM (Fig. 1F-H, Supplemental movie S5,6). All these results indicate the existence of vesicles directly transferred between adjacent cells. We named this phenomenon as Direct Intercellular Vesicle Exchange (DIVE) and these vesicles as InterCellular Vesicles (InterCVs).

### Kinetics of InterCVs

To clearly delineate cell boundaries while observing the behavior and dynamics of InterCVs, we conducted live imaging using CD81-SG-MDCK cells (green) and Lyn-Azalea-MDCK cells (red) seeded at a ratio of 1:100 (Fig. 2A). To accurately capture only the vesicles moving within and between cells, we trimmed and excluded the regions near the glass surface and the upper parts of the cells from the entire cell images, analyzing only the areas specifically at heights from 0.6 µm to 3.3 µm from the glass surface where the cells are present (Fig. S2A and S2B). Between the cell boundary of two green cells (box-i) in Fig. 2A), movements of green vesicles between these cells are often observed. Among them, an array of green vesicles was observed traversing from left to right cells across the cell boundary (Fig. 2B, arrowheads, supplemental movie S7), resembling observation shown in Fig. 1B. Vesicles traversing from green cells to red cells more clearly showed their kinetics (box-ii) in Fig. 2A). It was observed that green-labeled Vesicle1 moved from the right Green cell towards the left Red cell, paused briefly after reaching PM of Red cell, and then passed through PM to enter Red cell (Fig. 2C, supplemental movie S8). The time from contacting PM to passing through it was approximately 10 seconds, indicating highly rapid membrane passage. Moreover, even though some vesicles observed in Red cells are accompanied by red fluorescence, possibly incorporated with endocytosis (Fig. S2C), the green vesicle remained independently green after passing through PM without acquiring any red PM components from Red cell. These results suggest that this process is not mediated by endocytosis. Another green vesicle, not accompanied by red color (vesicle 2), was residing inside Red cell at the start of the observation. After briefly attaching to PM of Red cell, it moved away and relocated to almost the identical site of PM where vesicle 1 had passed. It then repeatedly moved across both sides of PM and into the interior of Green cell (Fig. 2C). During the repetitive crossing movement, vesicle 2 consistently passed through nearly the identical site on PM of Red cell. When the trajectories of vesicle 1 and vesicle 2 were compared, it became evident that both vesicles crossed almost the identical site on PM of Red cell (Fig. 2D). These observations suggest the presence of a specific route or structure, such as a pore, at the boundary between the two cells that allows vesicles to pass through. These vesicles are moving at approximately 0 to 1.0 µm/s, and the time required for the vesicles to cross PM after reaching the membrane was approximately 10 to 20 seconds (Fig. 2E). Other vesicles near the boundary between two cells moved in speed at less than 0.4 µm/s on average, with nearly 1.5 µm/s in maximum, and traveled approximately 10 to 40 µm in 4.4 min of observation period (Fig. 2F). We also evaluated the moving kinetics of CD81-positive vesicles overall in cells (Fig. S2D, supplemental movie S9). A two-dimensional live imaging captured at high speed (every 19 ms) allowed us to obtain images covering several cells almost identical to those observed under the microscope in real time. Tracking of CD81-positive vesicles (999 vesicles in total) demonstrated the presence of both actively moving vesicles and stationary ones (Fig. S2D and S2E). Vesicles defined as active by k-means clustering (top 117 cells in 999 cells) were observed moving at speeds exceeding 1.0 µm/sec on average (Fig. S2F). Thus, a large number of vesicles are actively moving within the cell, and at least some of them, as InterCVs, are directly transferred to adjacent cells through a previously unknown mechanism.

**Fig. 2.**
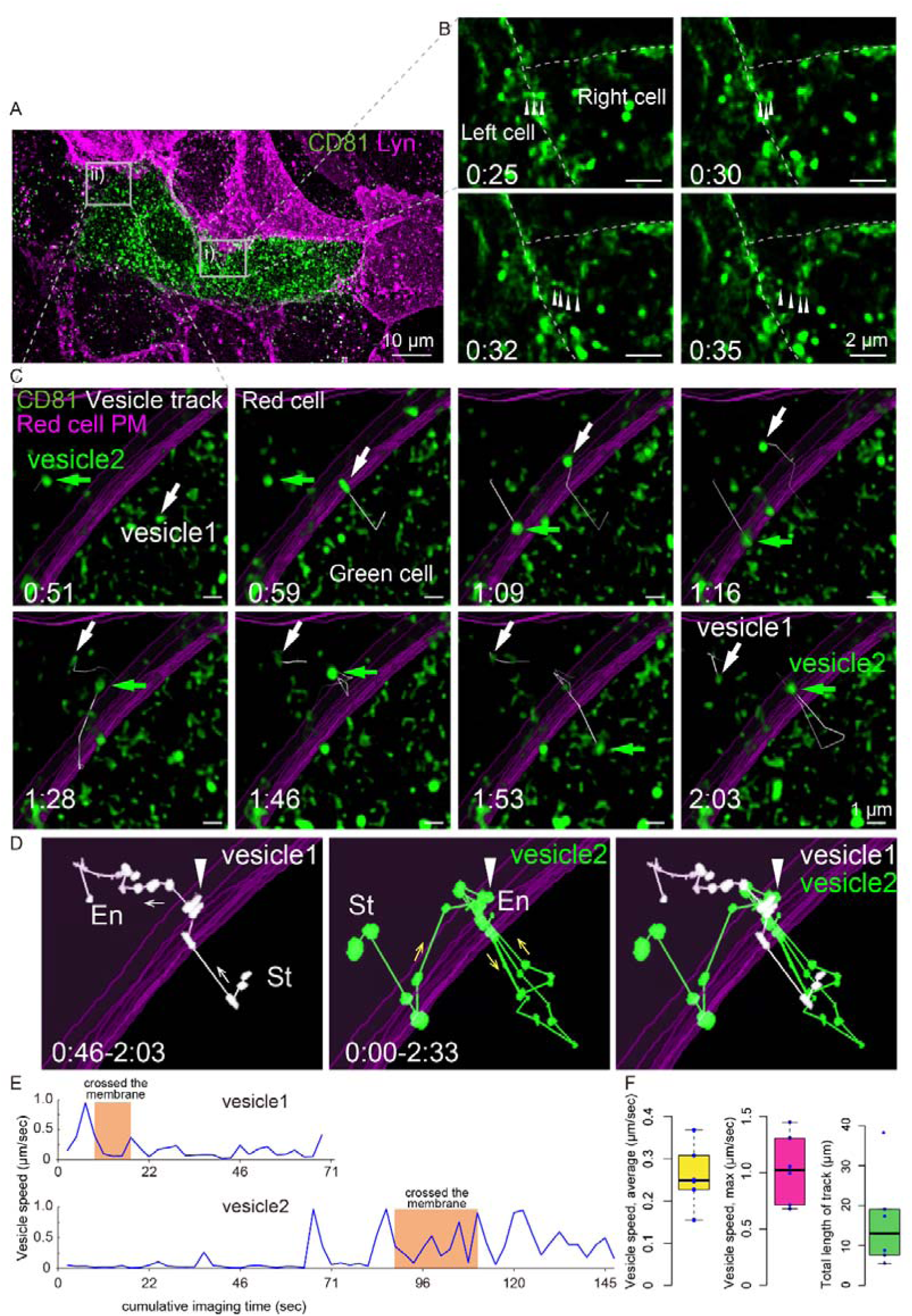
Kinetics of InterCVs captured by high spatiotemporal resolution live imaging. (**A**) Co-cultured CD81-SG-MDCK (green cells) and Lyn-Azalea-MDCK (red cells). Scale bar, 10 μm. (**B**) A row of vesicles traversing PM. Time series of the view (green color alone) within box (i) in (A) (captured from Supplemental movie S7). Time (min:sec), dotted lines: border between cells, arrowheads: vesicles. Note that an array of vesicles overlapping the border between cells (0:25) moves rightward into Right cells (0:30-0:35). Scale bars, 1 μm. (**C**) Vesicle transfer through a specific PM position. Time series of the view within box (ii) in (A) (captured from Supplemental movie S8). Magenta: plasma membrane of Red cell (left). Two vesicles of Green cell, vesicle 1 (white arrow) and vesicle 2 (green arrow), and their trajectories (white lines) are shown. Time (min:sec). Scale bars, 1 μm. (**D**) Trajectories of transferred vesicles. Overlaid images of vesicle positions in each frame (every 2.5 seconds) with their trajectories. Left panel: vesicle 1, middle panel: vesicle 2, right panel: overlay of vesicles 1 and 2. St, En: start and end points, respectively. Arrows: direction of vesicle movement. Arrowheads: positions of the vesicles crossing through PM. Note that PM crossing points of vesicle 1 and 2 are almost identical. (**E**) Speed of vesicles 1 and 2 during PM crossing. Vertical axis: speed. Horizontal axis: relative time points from the start of vesicle tracking. Orange squares: periods crossing PM. (**F**) Kinetics of vesicles moving near cell boundary. Average speed (left), maximum speed (center), and total distance traveled during the observation period of 4 minutes and 24 seconds (right). n=6.

### Ultrastructural analysis of InterCVs and DIVE with 3D electron microscopy

To identify structures that support the existence of InterCVs and DIVE mechanism, we explored and examined fine structures beyond the live imaging resolution using 3D electron microscopy analysis with FIB-SEM. CD81-SG-MDCK and Lyn-Azalea-MDCK were co-cultured, fixed, and examined with FIB-SEM. Fig. 3A-3C show an example of an array of vesicles looking like crossing PM as observed in Fig. 1B and 2B. The second left vesicle among four vesicles in the array is overlapping PM, suggesting that these vesicles should be InterCVs traversing PM. These vesicles are approximately 50 nm diameter and composed of single cell membrane. In another sample of a vesicle array at PM (Fig. 3D-3G), PM signal was partially unclear near the vesicle array (Fig. 3D, E). 3D reconstruction of each image showed that the unclear region may be able to form a pore-like structure located at the intersection of the vesicle array (Fig. 3F-3G). The other sample may show filaments and vesicles. For this sample, we employed CLEM (correlative light and electron microscopy) assay combined fluorescent imaging and FIB-SEM. We confirmed cell boundary area of CD81-SG-MDCK and mScarlet-H-MDCK cells in which many green vesicles were observed in red cells with fluorescent imaging, and then examined the area with FIB-SEM. In this sample, a number of periodical alignments of small black dots with 30-100 nm intervals across the cell boundary was observed (Fig. S3A and S3B). As shown in Fig. S3C and S3D, there were several vesicles along with the black dots array. Taken together with all these structural findings (Fig. 3 and S3) and InterCV kinetics in live imaging (Fig. 2), now we speculate the DIVE system as follows: PMs of adjacent cells fuse to form a pore-like structure, through which a kinds of filaments extend, allowing vesicles (InterCVs) to move fast along them by molecular motors and be exchanged between adjacent cells (Fig. 3H).

**Fig. 3.**
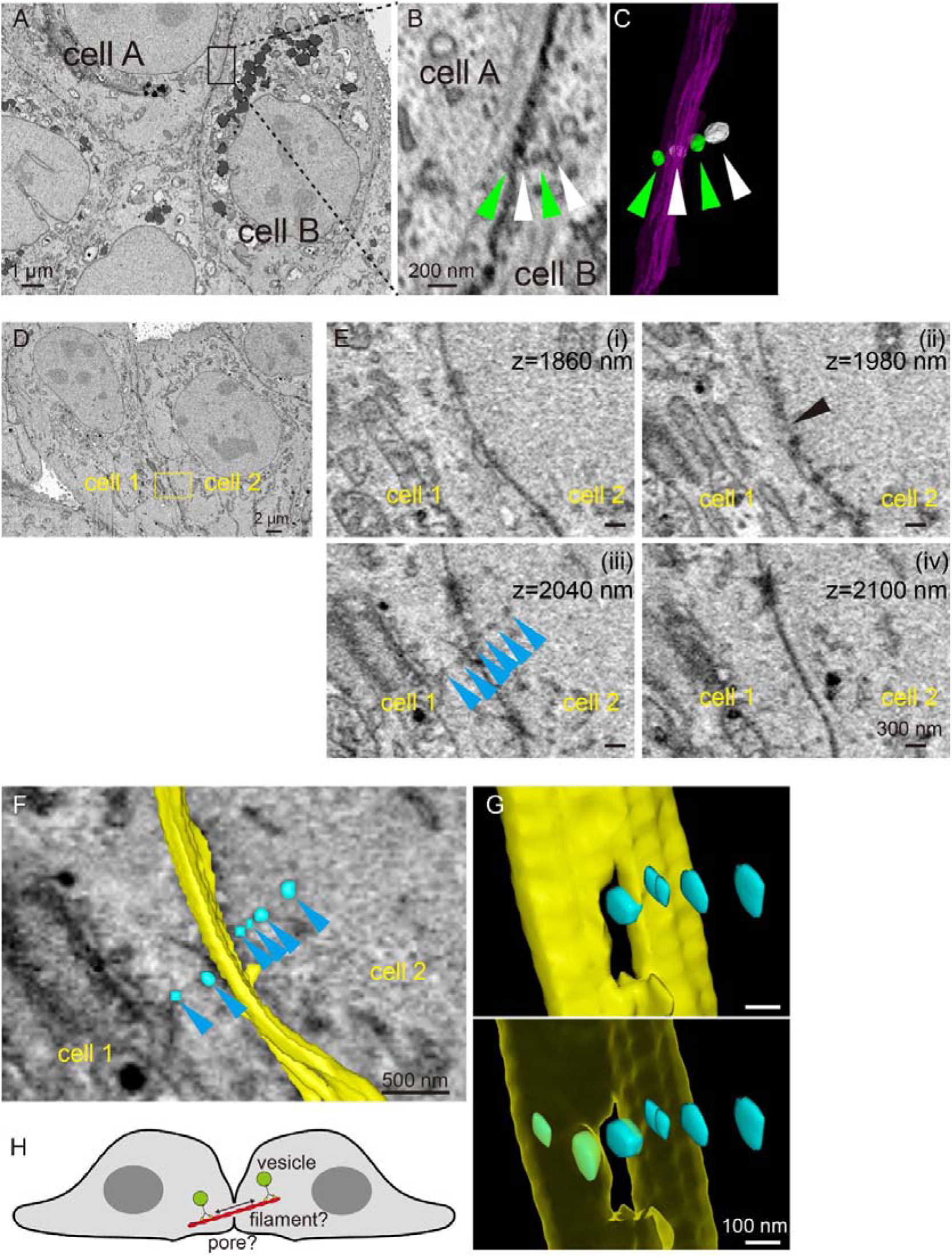
3D Electron microscopic observation supporting DIVE. (**A**-**C**) An array of vesicles traversing PM (**A**) FIB-SEM image of adjacent two MDCK cells (Cell A and Cell B). Scale bar, 1 μm. (**B**) Enlarged image of the black box in (A). Four vesicles forming an array across the boundary of Cells A and B. Vesicles are alternately indicated with green and white arrowheads for easy identification. Scale bar, 200 nm. (**C**) 3D image generated from each segment of the cell boundary surface (magenta) and vesicles (alternating white and green) from (B). The boundary surface shown as transparent. Note that the second left vesicle appears to overlap and traverse PM. (**D**-**H**) a pore-like structure for InterCV transfer (**D**) FIB-SEM image of adjacent MDCK cells (Cell 1 and Cell 2). Scale bar, 2 μm. (**E**) A magnified z-series of the yellow box area in (D). (i)(iv) PM are lined clearly. (iii) an array of vesicles with approximately 50 nm in diameter across PM (blue arrowheads). (ii) Unclear PM signal near the array of vesicles (arrowhead). Scale bars, 300 nm. (**F**) Overlay of the segmented cell boundary surfaces (yellow) and vesicles (cyan) with the original image. Scale bar, 500 nm. (**G**) 3D reconstructed Images (showing only the segmented cell boundary surface (yellow) and vesicles (cyan)). The cell boundary surface is shown as opaque in the upper panel and as transparent in the lower panel. Scale bars, 100 nm. Note that a pore-like structure intersecting the vesicle array is suggested. (**H**) Hypothetical schematic diagram of DIVE.

### Biological Function of DIVE

We previously reported that PSyC phenomenon was mediated by vesicles mainly between directly contacted cells. We confirmed the movement of CD81-positive vesicles containing miR-132 between mESCs (Fig. S4, supplemental movie S10). Then, it is considered that microRNA is a content of InterCVs and PSyC is one of the biological functions of DIVE. Recently, a report concerning intercellular mRNA transfer mainly through direct cellular contact between mESCs and human pluripotent stem cells (hPSCs) has been appeared [15]. ESCs can exist in two states: the naïve state, which resembles the epiblast of pre-implantation embryos and the primed state, which resembles the epiblast of post-implantation embryos [16, 17]. Mouse ESCs are basically in the naïve state whereas human PSCs are in the primed state. Yoneyama et al. reported that co-culture of mESCs and hPSCs induced the naïve conversion of hPSCs with intercellular mRNA transfer through direct contact of mESCs and hPSCs. Then, we confirmed the possibility that this intercellular mRNA transfer was mediated by DIVE mechanism. We co-cultured CD81-SG-mESC with human ESCs into which lifeact-mScarletH was introduced to mark actin fibers (lifeact-mScarletH-hESCs) (Fig. 4A). TFAP2C, a transcription factor reported to be important for the naïve state in hESCs [18] and one of the transferred mRNAs in Yoneyama’s study, was labeled with molecular beacons [19] targeting mouse TFAP2C mRNA. Mouse ESC-derived vesicles including mouse TFAP2C mRNA were observed to move back and forth between mouse ESCs and hESCs (Fig. 4B, C, Supplemental movie S11, 12). Mouse ESC-derived vesicles encapsuling mouse TFAP2C mRNA were found in hESCs moving along actin filaments (Fig. 4D) and some of them reached near hESC nucleus (Fig. 4E). These results suggest that intercellular mRNA transfer that contributes to naïve conversion of hPSCs should be mediated by DIVE with InterCVs containing mRNA, and exert function to induce PSyC to naïve conversion. In Fig. 1B, we observed InterCVs co-localized with DNA (DAPI). With all together, InterCVs should carry nucleic acids such as microRNA, mRNA, and DNA as cargo, and at least induce PSyC phenomenon among neighboring cells through DIVE mechanism. Finally, we preliminarily examined the molecular mechanisms and physiological roles of DIVE. We hypothesized that while mouse ESCs are cultured to maintain the naïve state, DIVE induces PSyC, leading to the synchronization of the naïve state and resulting in homogenous naïve cell appearance within each ESC colony. Then, mouse ESCs were cultured under naïve maintenance conditions and treated with various inhibitors such as exosome production inhibitors (nSMase2 inhibitor, Manumycin A), clathrin-dependent endocytosis inhibitor (chlorpromazine, CPZ), macropinocytosis inhibitor (5-(N-Ethyl-N-isopropyl) amiloride, EIPA), an inhibitor for macropinocytosis, phagocytosis, and actin polymerization (Cytochalasin B), and actin polymerization inhibitor (Chaetoglobusin A). We evaluated naïve cell homogeneity with uniformity of Nanog expression levels in each cell within the colony. Though all inhibitors significantly reduced the uniformity of Nanog expression levels, actin polymerization inhibitor (Chaetoglobusin A) showed significantly larger differences and mosaic pattern in Nanog expressions than other inhibitors (Fig. 4F, G), suggesting that actin polymerization is one of the components of DIVE mechanism. To examine the physiological roles of DIVE in vivo, we performed ex vivo culture of mouse blastocysts. The blastocyst at embryonic day 3.5 contains an inner cell mass (ICM) within its cavity, which is divided into two groups of cells: Nanog-positive epiblast and Gata4-positive primitive endoderm, each forming distinct layer. Appearance of isolated Gata4-positive cells ectopically from ICM region is rare event in normal culture condition (isolated; Fig. 4H). The addition of Manumycin A apparently increased the frequency of embryos with isolated-type Gata4 expression (Fig. 4I), suggesting that DIVE may induce PSyC for Gata4-positive primitive endoderm formation in ICM. Thus, DIVE should play a physiological role even in vivo conditions at least such as embryogenesis.

**Fig. 4.**
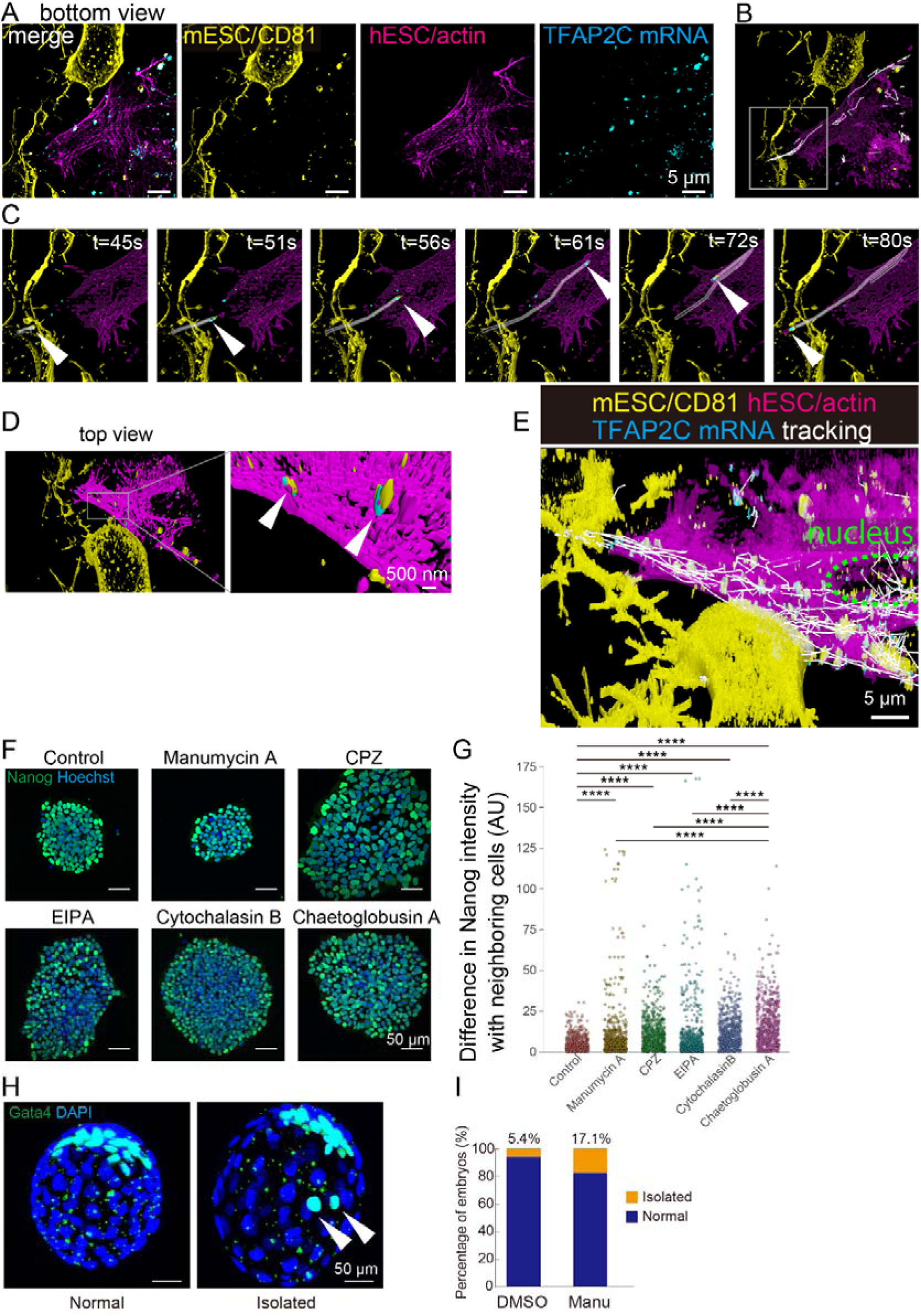
Putative functions and roles of DIVE in PSyC. (**A** – **E**) InterCV transfer between adjacent mouse and human pluripotent stem cells (roles in naïve conversion). (**A**) 3D image of co-cultured CD81-SG-mESCs (yellow) and lifeact-mScarletH-hESCs (magenta). mRNA of mouse-derived TFAP2C was labeled with molecular beacons (blue). Scale bars, 5 μm. (**B** and **C**) Transfer of mouse InterCVs with mouse mRNA to human cells. (**B**) Segment view showing the trajectories of mouse-derived vesicles carrying TFAP2C mRNA (**C**) A time series within the white box area in (**B**) (Captured from Supplemental movie S11,12). White arrowheads: a mouse-derived vesicle (yellow) with mouse mRNA (blue) moving back and forth between mESC and hESC. (**D**) A top view of the segment view (a flipped image from (**A**)). Right panel: an enlarged view of the box in Left panel. White arrowheads: mouse-derived vesicles (yellow) encapsuling mouse TFAP2C mRNA (blue) on actin filaments in hESCs (magenta). Scale bar, 500 nm. (**E**) Overlaid time series images of (**D**). Green dashed line: Nuclear region of hESC. Note that a number of mouse vesicles (yellow), some of which including mouse mRNA (blue), reached near or in adjacent human cell nucleus. Scale bar, 5 μm. (**F** and **G**) PSyC for homogeneity of pluripotent stem cell colony. (**F**) Immunostaining for Nanog, an undifferentiation marker, in mouse ESC colony. Representative stained ESC colonies are shown. Nanog (green), DAPI (blue). mouse ESC colonies were treated with indicated reagents. Scale bars, 50 μm. (**G**) Heterogeneity of Nanog expression among each cell in colonies. Plot of the absolute difference in Nanog intensity between each cell and its three closest neighbors, with each group comprising N=586 (586 neighbors). **** p ≤ 0.001. (**H** and **I**) PSyC in embryo development (**H**) Immunostaining for Gata4, a primitive endoderm marker, in mouse embryos corresponding to E4.5 (one day after the ex vivo culture of E3.5 embryos). Gata4 (Light blue) and DAPI (blue). Representative normal (left) and isolated type (right) Gata4 expression. Normal: Gata4-positive cells exclusively in ICM area. Isolated: isolated ectopic appearance of Gata4-positive cells (arrowheads). Scale bars, 50 μm. (**I**) Quantification of the proportions of normal and isolated patterns in mouse embryos treated with DMSO (vehicle, n=37) or Manumycin A (Manu, n=35).

## Discussion

In this study, we discovered a novel modality of intercellular communication, mediated by direct vesicle transfer across PM between adjacent cells, which we named DIVE. By employing high spatiotemporal resolution live imaging, we were able to detect rapidly moving vesicles and observed that vesicles, 50–500 nm in diameter, sometimes moving in an array, traversed specific sites on PM repeatedly, and crossed PM rapidly (within 10-20 seconds). These rapid movement kinetics and the lack of co-localization of vesicles with the recipient cell PM components after crossing PM suggest that this process is not mediated by endocytosis. Furthermore, no specialized structures such as protrusions, commonly observed with cytonemes and TNTs, were detected, indicating that this is a previously unrecognized modality of vesicle transfer. The observed features of DIVE now lead us to hypothesize a model shown in Fig. 3H. Though observations with 3D electron microscopy analysis (Fig. 3 and S3) can support our proposed model, further optimization of conditions is necessary to obtain definitive evidence. Our current model remains a hypothesis that requires further detailed and precise investigation into the nature of DIVE.

The discovery of this previously unidentified phenomenon, DIVE, was made possible primarily due to advances in live imaging techniques. The observed InterCVs are not only small in size but also exhibit unexpectedly rapid movement, making it difficult to capture their trajectories. By combining the use of StayGold, a novel fluorescent probe with high brightness and high photostability properties, with high-resolution microscopy, we were able to track vesicle movements, particularly those that cross PM within a short period, with a temporal resolution of approximately 2 seconds across a space of 128x128x2.7 µm. Recent attention has focused on protrusion-based structures, such as TNTs and cytonemes, for their roles in intercellular vesicle transfer. These structures are visible under microscopy or electron microscopy, and the movement of vesicles within these tubular structures can be detected using conventional spatiotemporal resolution. However, the existence of DIVE and InterCVs necessitated the level of spatiotemporal resolution we demonstrated here for their detection. In addition, our previous discovery of PSyC revealed that PSyC arises primarily through the exchange of vesicles between adjacent cells, providing strong motivation to observe vesicle transfer between cells in real time. This convergence of research motivation and advances in imaging technology enabled the discovery of DIVE.

Investigating the structure underlying DIVE remains challenging even with 3D electron microscopy, as observations are limited to fixed samples, making it difficult to determine whether detected vesicles are in the process of crossing the membrane. Identifying the molecular components involved in DIVE and InterCV formation and verifying their localization through both live imaging and ultrastructural analysis are necessary steps. Potential components involved in DIVE include molecules responsible for membrane fusion and pore formation, filaments connecting cells, molecular motors, and others contained within InterCVs. InterCVs confirmed by electron microscopy are structures with a single membrane, with diameters of 50-100 nm observed in electron microscopy and approximately 100-500 nm in live imaging. Candidates of previously reported single-membrane vesicles include naked EVs, endosomes, Golgi vesicles, lysosomes, and amphisomes [20]. InterCVs were marked by CD63 and CD81 in this study, suggesting they may share a biogenesis pathway with EVs or endosomes. The cargo carried by InterCVs includes nucleic acids— DNA, mRNA, and microRNA—but other potential cargoes remain to be explored. Similar to EVs, a variety of proteins may also be included. Although we did not observe mitochondrial transfer in our study, the possibility of mitochondrial transport, as seen in TNTs and EVs, cannot be ruled out. The observation of relatively large vesicles suggests the possibility of broader intercellular organelle exchange, which warrants further investigation.

At present, PSyC is considered to be one of the candidate biological functions of DIVE. Our previous findings identified miR-132 as a molecule that induces synchronization of ESC differentiation, and its transfer between adjacent cells via vesicles was visualized. The results presented in Fig. 4 further support the role of DIVE in PSyC. While PSyC itself is a relatively new discovery, requiring further accumulation of biological data, the synchronization of cell behavior within cell clusters is speculated to be critical for processes such as multicellular organism formation, embryonic development, and organ formation and function. Elucidating the involvement of PSyC and DIVE in these processes would significantly enhance their biological importance.

The potential implications of DIVE extend beyond current understanding. PSyC-driven cellular phenotype control and/or targeted vesicle exchange between cells could pave the way for the development of novel therapeutic strategies. These strategies may include cancer treatment and anti-aging therapies through the introduction of designed cells into the human body. Thus, the discovery of DIVE reveals a previously unrecognized modality of intercellular communication that may lead to a paradigm shift in biology, reexamining cellular interactions and potentially inspiring innovative biomedical applications. This research carries significant transformative potential for biology and medicine.

## Limitation

The data presented in this study primarily focus on the observation and detection of the novel phenomenon through live imaging. Further details, particularly regarding its molecular mechanisms, remain to be elucidated. The model of the DIVE mechanism proposed here is based on observations of the phenomenon and will require verification and refinement through various research approaches in the future. Additionally, the biological significance of the DIVE phenomenon itself needs to be further explored and better understood in future studies.

Technically, high spatiotemporal resolution imaging like that used in this study (3D images consisting of several dozens of optical sections at 200-300 nm intervals, captured every 2-3 seconds) is particularly challenging for long-term imaging. The current limit is about 200 frames, or 6-10 minutes in total. This limitation is due to software and hardware processing constraints, including the explosive increase in data size and the fading of probes other than SG. It is anticipated that these barriers will be overcome with the development of advanced hardware, software, and probes capable of surmounting these limitations.

## ACKNOWLEDGEMENTS

We thank Dr. Masahide Kikkawa and Dr. Chieko Saito [Graduate School of Medicine, The University of Tokyo, Bunkyo-ku, Tokyo] for their invaluable technical and scientific support in FIB-SEM imaging using JIB-PS500i.

We thank Dr. Takahiro Maeda [TAKARA Bio, Shiga, Japan] for editing plasmids. Live-cell imaging with super-resolution and light sheet microscopies was performed at ZEISS-iCeMS Innovation Core of iCeMS Analysis Center, Kyoto University.

## Author contributions

T.M. and J.K.Y. conceived and designed the whole study. T.M. performed all the experiments and analyzed the data. T. M. and J.K.Y. wrote the manuscript. T.K. and K.O.F. performed FIB-SEM imaging using a Crossbeam 540. R.A. and A.M. provided new materials and technically supported imaging studies. F.I. and T.K.F. assisted with light sheet imaging and super resolution imaging. D.C. cultured and prepared mouse embryo samples. S.T. and Y.T designed the molecular beacons and prepared cationized gelatin spheres. Y.S. provided new material and methods. J.K.Y. conceived the project, secured funding.

---

**Fig. S1.**
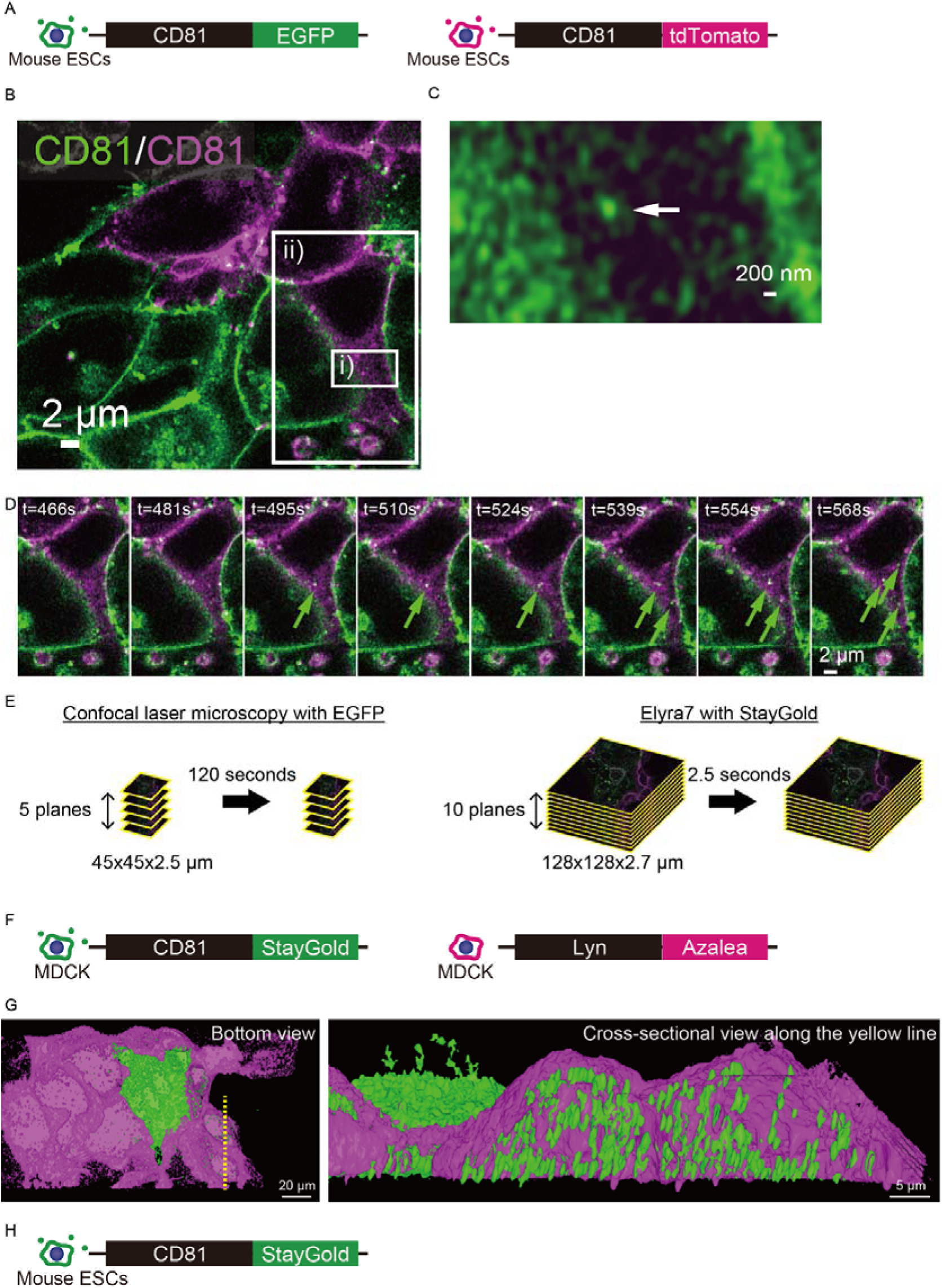
Imaging with conventional FP and improvement with StayGold. (**A**) Mouse ESCs carrying CD81-EGFP fusion protein (CD81-EGFP-ESC) and carrying CD81-tdTomato fusion protein (CD81-tdTomato-ESC). (**B**) Co-culture of CD81-EGFP-ESC and CD81-tdTomato-ESC. CD81-FP fusion proteins mark cellular membranes including vesicles and plasma membrane. Scale bar, 2 μm. (**C**) Magnified view of the white box i) in (B). White arrows indicate CD81- positive vesicles of approximately 200 nm in diameter. Scale bar, 200 nm. (**D**) Moving green vesicles in red cells. Time series of the magnified view from white box ii) in (B) (captured from Supplemental movie S1). Green arrows: green vesicles moving in a red cell. Scale bar, 2 μm. (**E**) Improvement of spatiotemporal resolution. Left: Schematic diagram of 4D imaging using EGFP and confocal fluorescence microscope LSM880. Right: Schematic diagram of 4D imaging using StayGold and Elyra7. (**F**) MDCK cells carrying CD81-StayGold fusion protein (CD81-SG-MDCK) and carrying Lyn-Azalea fusion protein (Lyn-Azalea-MDCK). (**G**) Confocal microscopic confirmation of green vesicles in adjacent red cell. Left: Segmented bottom view of co-cultured CD81-SG-MDCK (green cells) and Lyn-Azalea-MDCK (red cells), Right: Cross-sectional view along the yellow line in left panel. Many green vesicles are in the red cell (to facilitate the visualization of vesicle positions, the Z-axis is displayed at twice its original scale). Scale bars, 20 μm and 5 μm, respectively. (**H**) Mouse ESCs carrying CD81-StayGold fusion protein (CD81-SG-mESC).

**Fig. S2.**
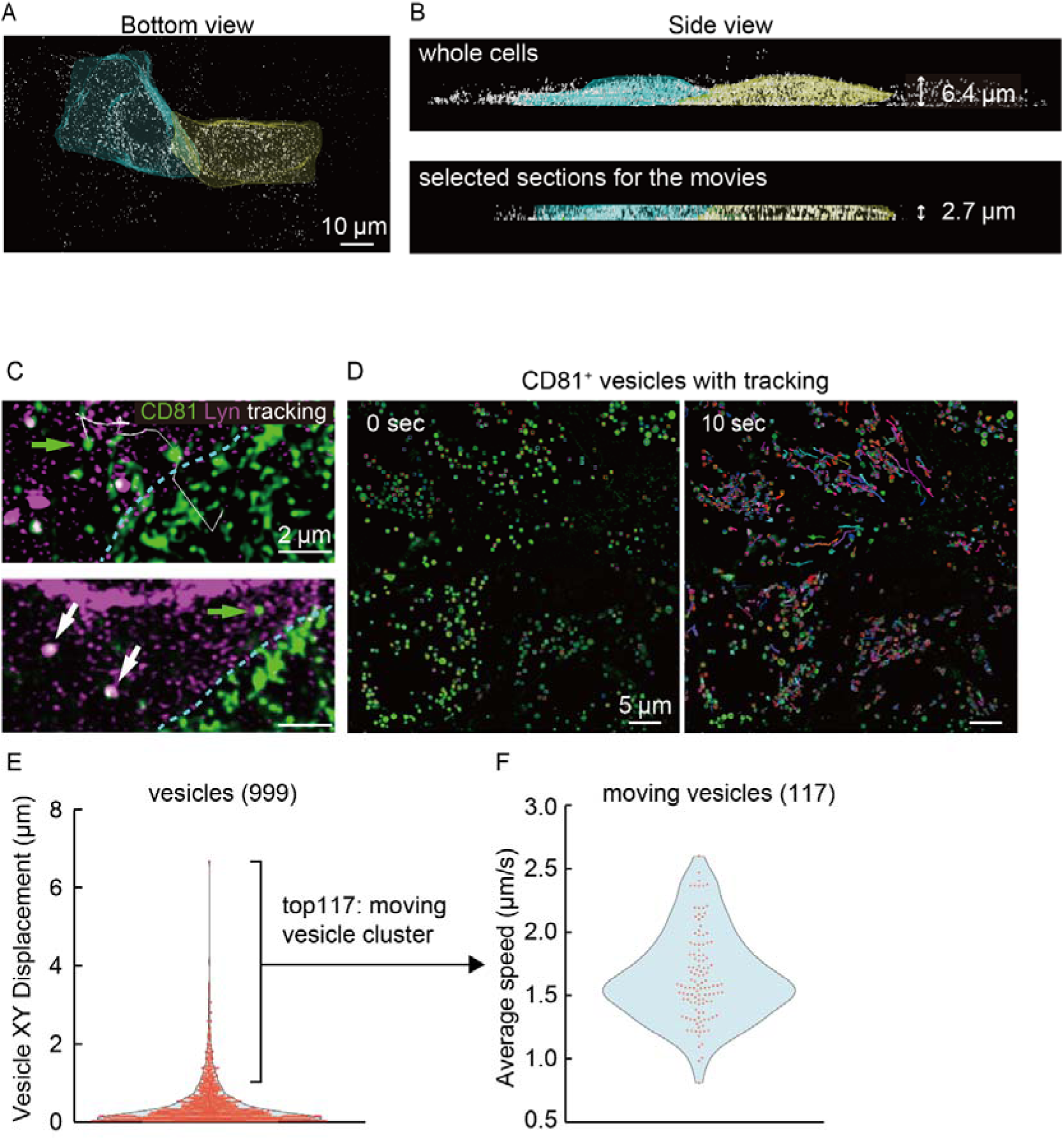
Analysis of moving vesicles. (**A**) Bottom view of the two green cells from Fig. 2A (segmented image). The left cell is shown in cyan, and the right cell in yellow. Scale bar, 10 μm. (**B**) Side view of the two cells. Upper panel: the entire cells. Lower panel: selected area after trimming of near glass surface and upper parts of cells. The area selected for analysis was within the interior of the cells. (**C**) Transferred vesicles with or without PM components. Boundary region between Red and Green cells (Fig. 2). Cyan dashed line: cell boundary. Upper panel, Green arrow: a green vesicle that have moved from Green cell to Red cell. Lower panel, the same field after time has elapsed. Green arrow: the green vesicle moved from Red cell still independent (not enclosed by Lyn). White arrows: green vesicles enclosed by Lyn (white, double positive for green and magenta), suggesting vesicles transferred via endocytosis. Scale bars, 2 μm. (**D**-**F**) Real-time vesicle movement under microscopy (**D**) 2D live imaging of CD81-SG-MDCK cells with tracking applied to CD81- positive vesicles. Left panel: the start of imaging (time 0). Right panel: 10 seconds later. Vesicles in several cells are tracked. Scale bars, 5 μm. (**E**) XY displacement of vesicles (999 cells in total) from the data in (D). (**F**) Plot of speeds of vesicles defined as “moving vesicles” (117 cells) after dividing G into two groups using k-means clustering.

**Fig. S3.**
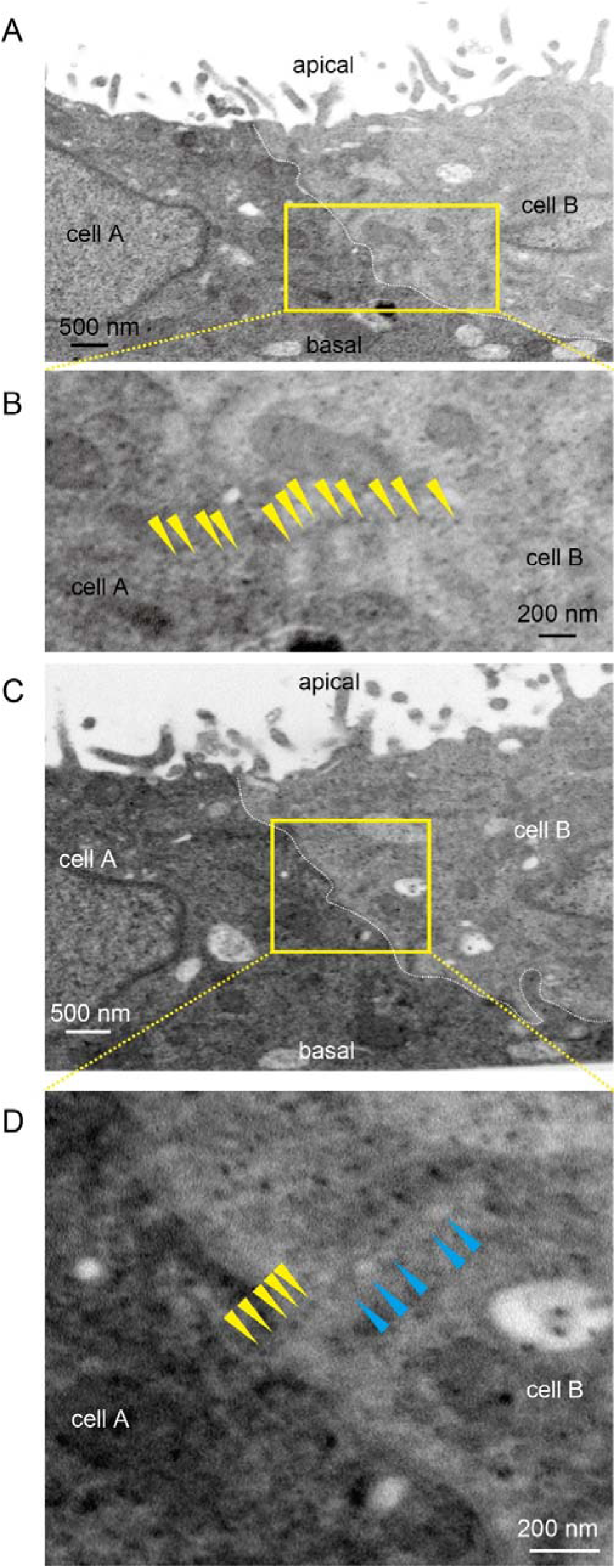
Arrays of black dots across the cell boundary. (**A** and **B**) Periodic black dots across the cell boundary. (**A**) A cross-sectional FIB-SEM image of cell boundary area between cell A and cell B. (**B**) Enlarged view of the yellow box in (A). Yellow arrowheads: an array of black dots extending from cell A to cell B region across the cell boundary. (**C** and **D**) Co-localization of arrays of vesicles and black dots at cell boundary. (**C**) A cross-sectional FIB-SEM image of cell boundary area between two MDCK cells, cell A and cell B (different cross-section from (A)). White dashed line: the boundary between cell A and cell B. Scar bar: 500 nm. (**D**) Enlarged view of the yellow box in (C). Yellow arrowheads: an array of black dots. Light blue arrowheads: an array of vesicle-like structures.

**Fig. S4.**
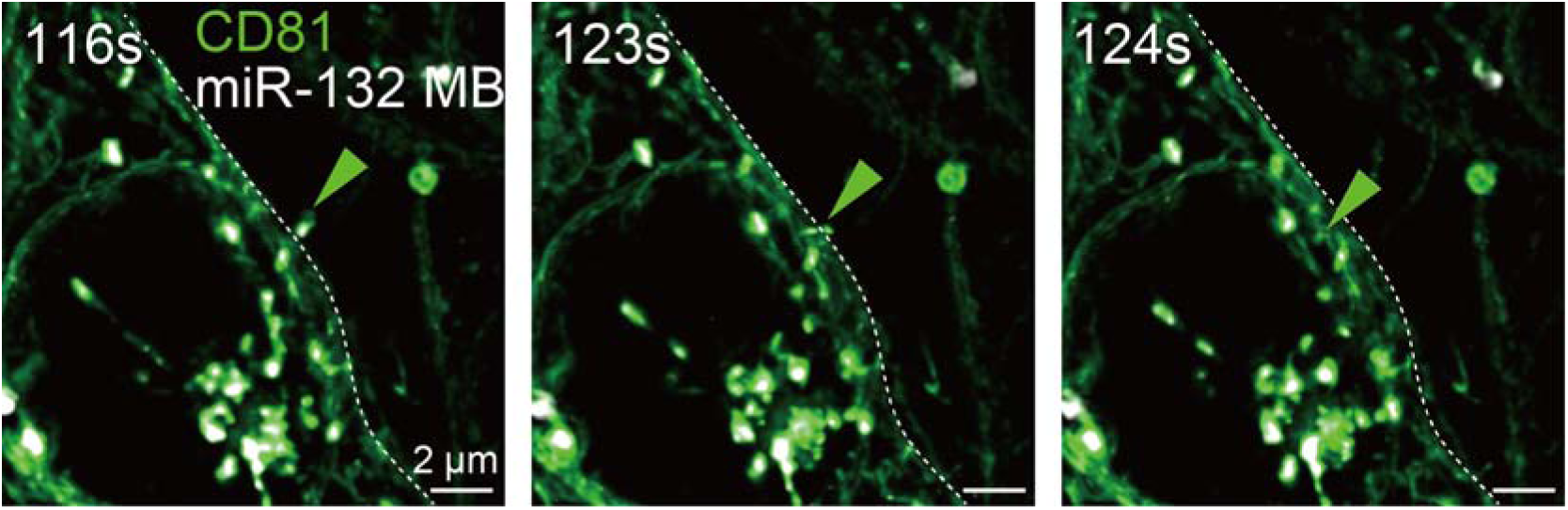
A vesicle containing miRNA crossing the cell boundary. A Time series of 3D live imaging at a cell boundary of CD81-SG-mESCs (captured from Supplemental movie S10). Green vesicles: CD81-SG-mESC-derived vesicles. White: miR-132 marked with molecular beacon (MB). White dashed line: the boundary between two cells. Green arrowheads: a vesicle containing miR-132 moving from right to left cell. Scale bar: 2 μm.

## Materials and methods

### Cell lines

Information about the cell lines used and the plasmids introduced into the cells in each experiment is summarized in Table 1.

**Table1:**
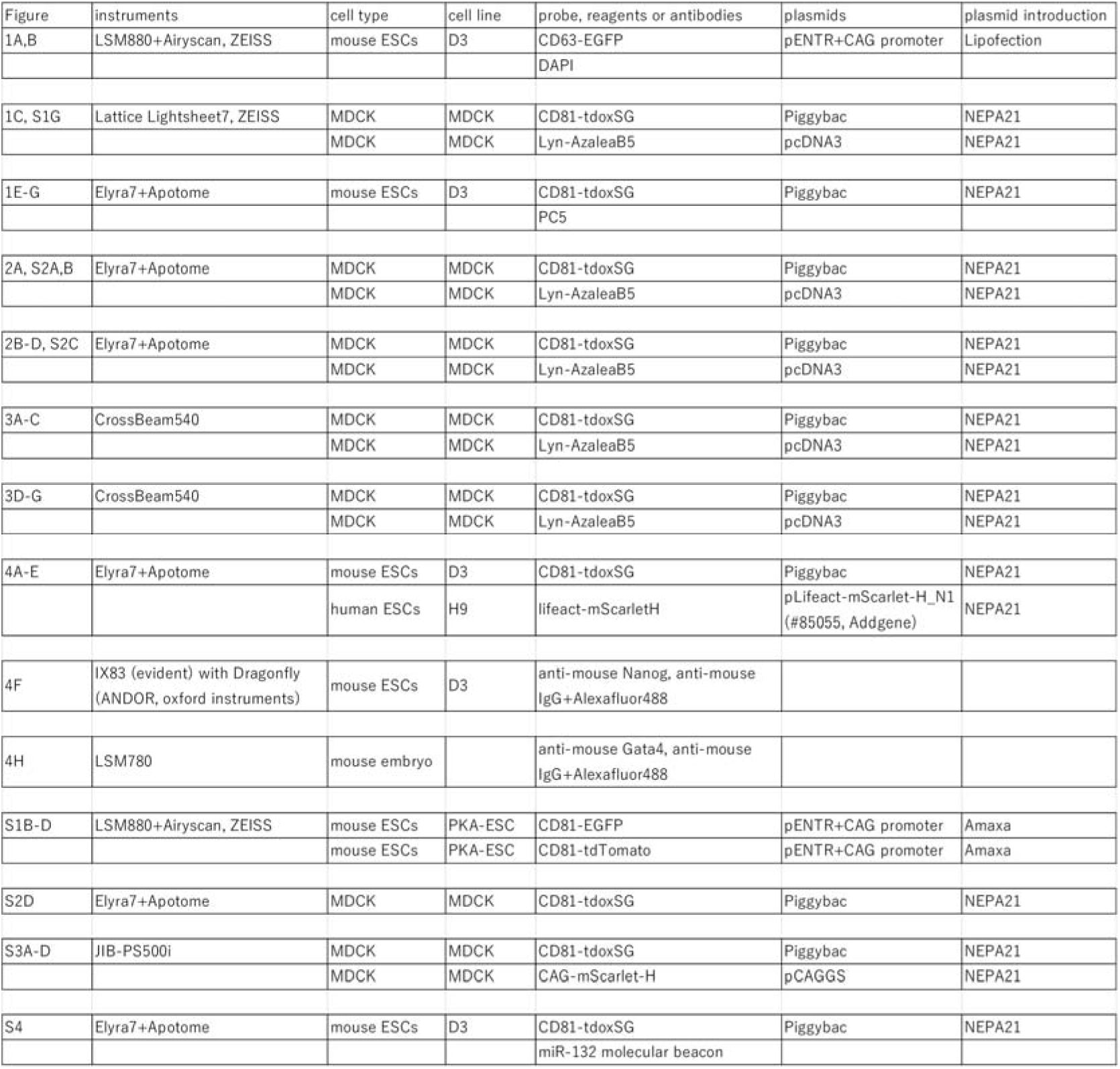
Instruments and fluorescent probes used for data acquisition in each figure.

ES-D3: A mouse ESC line derived from 129/Sv blastocysts [21].

PKA-ESC: A mouse ESC line carrying a tetracycline-regulatable constitutively active form of the PKA (CA-PKA) gene [22]. It was generated as described previously. The PKA-ESC line behaves similarly to the wild type when doxycycline (DOX) is added. ESC lines were maintained in Glasgow Minimum Essential Medium (GMEM; 11710-035, Thermo Fisher Scientific, USA) containing 10% KnockOut Serum Replacement (KSR; 10828-028, Thermo Fisher Scientific), 1% fetal bovine serum (FBS; SAFC Biosciences, USA), 2,000 units/mL leukemia inhibitory factor (LIF; Merck Millipore), 1 mM sodium pyruvate (Sigma-Aldrich, Merck KGaA, Germany), 1% MEM Non-Essential Amino Acids (NEAA; 11140-050, Thermo Fisher Scientific), 50 U/mL penicillin (Meiji Seika Pharma, Japan), 50 µg/mL streptomycin (Meiji Seika Pharma, Japan) and 0.1 mM 2-mercaptoethanol (2-ME; 21985-023, Thermo Fisher Scientific). The culture dish was pre-coated with 0.1% gelatin at 23 °C for 5min. For the maintenance of PKA-ESCs, 1 µg/mL Dox was added.

H9: The human ESC line developed by Thomson et al [23], kind gift from Dr. Thomson. H9 cells were maintained in StemFit (AK02N, Ajinomoto Healthy Supply, Japan). The culture dish was pre-coated with a solution prepared by mixing Matrigel (356231, Corning) and knockout DMEM (10829-018, Thermo Fisher Scientific) at a ratio of 1:60 and incubating at 37 °C for 1 hour.

MDCK II (Madin-Darby Canine Kidny Strain II) :An epithelial-like cell line, isolated from dog kidney, was purchased from DS Pharma Biomedical (Japan). MDCK cells were maintained in Dulbecco’s Modified Eagle Medium:Nutrient Mixture F-12 (DMEM/F12 (1:1); 11330-032, Thermo Fisher Scientific] containing FBS, 1%NEAA, 50 U/mL penicillin and 50 µg/mL streptomycin. The culture dish was pre-coated with 0.1% gelatin at 23 °C for 5min.

Plasmids were introduced into the cells by either NEPA21 typeII (NEPAGENE, Japan), Amaxa Mouse ES Cells Nucleofector Kit (VPH-1001, Lonza, Switzerland) or Lipofectamine LTX and Plus Reagent (15338-100, Invitrogen, Thermo Fisher Scientific, USA).

The settings for NEPA21 are as follows:

Porating Pulse: Voltage 150V, pulse width 5 ms, pulse interval 50 ms, number of pulses 2, attenuation rate 10%, polarity +

Transfer Pulse: Voltage 20V, pulse width 50 ms, pulse interval 50 ms, number of pulses 5, attenuation rate 40%, polarity +/-

### Plasmid construction

For each experiment, the plasmids used are listed in the table1. CAG promoter sequence was inserted between SalI and EcoRI sites on the multicloning site of pENTR3C (A10464, Thermo Fisher Scientific) using Ligation high Ver.2 (LGK-201, TOYOBO, Japan). The coding sequence (CDS) of CD63 or CD81 were trimmed of their stop codon and were fused with fluorescent proteins (FPs) such as EGFP, tdTomato, tdoxStayGold (tdoxSG) [12]. Each sequence was cloned using KOD Plus NEO (KOD-401, TOYOBO) or KOD One (KMM-201, TOYOBO), and inserted into pENTR3C either with or without CAG promoter using Ligation high Ver.2, In-Fusion HD Cloning Kit (TaKaRa Bio, Japan), or NEBuilder HiFi DNA Assembly Master Mix (#E2621S, New England Biolabs, NEB, USA). The CD63/CD81-FPs sequences were then transferred from the pENTR3C plasmid into the piggyBac vector [24] pPV EF1a-GW-iPA using Gateway LR Clonase II Enzyme Mix (11791-020, Thermo Fisher Scientific).

### Co-culture of mouse ESCs and human ESCs

ES-D3 were pre-cultured for several passages under naïve maintenance conditions (ESC maintenance medium supplemented with 3 µM of CHIR and 1 µM PD03) [25]. One day prior to the start of imaging, ES-D3 and H1 cells were mixed at a 1:1 ratio and seeded onto a matrigel-coated 8-well chamber and cultured in StemFit medium (AK02N, Ajinomoto, Japan).

### mRNA tracking

The mRNA of TFAP2C was tracked using a molecular beacon (MB) [Integrated DNA Technologies (IDT), USA] The sequence of MB was 5’-[Alexa647]-CGCGATC-TTTCAAGAGAGGGAACGGAT-GATCGCG-[Iowa Black RQ]-3’.

CGCGATC and GATCGCG form a stem structure, TTTCAAGAGAGGGAACGGAT is an mRNA recognition sequence, and Iowa Black RQ is a quencher suitable for Alexa647. Cationic gelatin nanospheres were employed for efficient delivery and prolonged release of molecular beacons into cells [19].

### Live-cell imaging

Live imaging was performed using several systems at 37 °C under 5% CO2. The LSM880+Airyscan (Carl Zeiss) was paired with a Plan-Apochromat 40x/1.4 Oil DIC M27 objective, with images processed via Airyscan using Zen 3.0 (black edition, Carl Zeiss) software. For the Elyra7 with Lattice SIM or Apotome (Carl Zeiss), the same objective was utilized, and images were processed with SIM2 using Zen 3.0 (black edition) software. The Lattice LightSheet7 (Carl Zeiss) used a 44.83x/1.0 NA objective, while the Dragonfly200 (Oxford Instruments Andor, UK) was combined with an IX-83 (Evident, Japan), and a UPlanXApo 100x/1.45 Oil objective was employed. For nuclear labeling, DAPI (4’,6- diamidino-2-phenylindole, 50 ng/mL) or PC5 (1 µM) was added to the culture medium. PC5 dye is now commercially available (Kakshine Red: KA-050R, Cosmo Bio, Japan). The excitation wavelength (Ex) and emission range (Em) for each fluorescent probe are as follows: DAPI (405/420-480, 495-550 nm), EGFP (488/420-480, 495-550 nm), SG (488/505–545 nm) for Lattice LightSheet7 and (488/420-480, 495-550 nm) for Elyra7, AzaleaB5 (561/570-620, 655- nm), PC5 (642/570-620, 655- nm), mScarlet-H (561/570-620, 655- nm), tdTomato (561/495-550, 570- nm), MB (642/570-620, 655- nm).

### Analysis of Nanog expression intensity in ESC colonies

ESC colonies were imaged using the Dragonfly system, with cells in a single layer close to the glass surface used for analysis. The xy coordinates of the center of each cell nucleus and the average Nanog intensity across the entire nucleus were outputted by arivis Vision4D 4.1.0 (arivis, Germany). The absolute difference in Nanog intensity between any given cell and its three nearest neighbors (only those within a distance of less than 20 μm) was calculated. To equalize the sample size, data for 586 cells were randomly sampled in each group. Statistical analysis was performed to determine if there were significant differences in the mean values among the control and various drug-treated groups. Only statistically significant differences are shown. These calculations, statistical analyses, and plots were conducted using R version 4.3.0 and associated packages (readxl, dplyr, tidyr, writexl, ggplot2, stats).

### Vesicle count

After performing segmentation of vesicles and cell regions separately using arivis Vision4D, the number of vesicles present in the green cells, red cells adjacent to green cells, and cells two cells away from green cells, was counted using the ‘Compartments’ operation.

### Focused ion beam scanning electron microscope (FIB-SEM) tomography

For the FIB-SEM imaging with CrossBeam540 (Carl Zeiss), resin embedded samples were prepared with slight modifications to the previously reported method [26]. In short, cells were fixed in prewarmed 4% paraformaldehyde (PFA) solution for 1 hour at 23°C, then transferred to a solution containing 2% glutaraldehyde (GA) 4% PFA solution with 0.1% tannic acid, and incubated for 12 hours at 4°C. The cells were then washed twice with 2% GA in 4% PFA solution for 10 minutes at 23°C. The samples were post-fixed for 2 hours at 4°C in deionized water (DW) containing 1.5% potassium ferrocyanide and 2% osmium tetroxide. After washing with distilled water (DW), the specimens were immersed in 2% osmium tetroxide in distilled water and washed with distilled water. The specimens were subjected to en bloc staining overnight in a solution of 1% uranyl acetate dissolved in DW to enhance contrast, followed by another wash in DW. Subsequently, the specimens were further stained with Walton’s lead aspartate solution for 30 min at 60°C. After the final DW wash, the samples were dehydrated through a graded ethanol series (50%, 60%, 70%, 80%, 90%, 99%, 100%) and embedded in Epon812, polymerized 40C for 12 hrs and 60C for 48 hrs. For the FIB-SEM imaging with JIB-PS500i (JEOL, Japan), correlation light and electron microscopy (CLEM) method was applied to record the position of the cell-cell boundaries in cross-sectional view and to target the location of the region of interest. CLEM sample preparation was performed as previously described [27], other than 0.1 M phosphate buffer (pH 7.2) was used instead of sodium cacodylate buffer, and osmification (performed with 2% Osmium tetroxide solution in 0.1M phosphate buffer without potassium ferrocyanide).

The specimens were set on the stage of a FIB-SEM (CrossBeam 540 or JIB-PS500i). In microscopy with CrossBeam540, Smart FIB software (Carl Zeiss) was used to mill the surface of the specimen with a gallium ion FIB. The SEM images were obtained by 370 or 156 times of the slice-and-view processes, so that the total sliced thickness was 3.7 or 4.6 μm. The resultant image stacks were processed and edited with Dragonfly 2022 software (ORS, Canada) In microscopy with JIB-PS500i, A carbon deposition layer was created on the upper surface of the observation target area. Subsequently, a trench was formed in front of the observation target area using a gallium ion FIB at 30 kV with a current of 30 nA. Following this, in order to smooth the surface of the trench wall that would serve as the observation face, the area was milled in two steps using a gallium ion FIB at 30 kV with currents of 10 nA and 3 nA. During imaging, milling was performed using a gallium ion FIB at 30 kV with a current of 3 nA, at a pitch of 10 nm per slice, and SEM images were obtained at a landing energy of 3 kV with Ultra High Resolution mode with RBED detector. The SEM images were obtained by 454 times of the slice-and-view processes, so that the total sliced thickness was 4.5 μm. The resultant image stacks were processed and edited with Fiji [28].

Segmentation of the plasma membrane and vesicles, as well as 3D reconstruction, were performed using arivis Vision 4D.

### Superovulation, embryo collection, and embryo culture

Eight- to ten-week-old ICR female mice were superovulated by injecting 5 IU of equine chorionic gonadotropin (eCG; ASKA, Japan), followed by 5 IU of human chorionic gonadotropin (hCG; ASKA, Japan) 48 hours later. Subsequently, the female mice were naturally mated with male mice overnight, and 1-cell embryos were collected the following day. The collected embryos were washed three times in KSOM supplemented with amino acids [29] and 4 mg/ml BSA, and then were cultured until embryonic day 4.5 (E4.5) in KSOM drops under mineral oil (Sigma-Aldrich) at 37°C in an atmosphere of 5% CO2. For the manumycin A treatment experiments, the embryos were transferred to KSOM drops containing manumycin A (7 µM) on E3.5.

### Immunofluorescence staining of embryos

Embryos were fixed with 4% paraformaldehyde at 4°C for 20 minutes and washed three times with PBS containing 0.3% polyvinylpyrrolidone (PBS/PVP, Nacalai tesque, Japan). Embryos were then permeabilized for 40 minutes with 0.5% Triton X-100 (Sigma Aldrich) in PBS. After being washed three times with PBS/PVP, the embryos were transferred to a blocking buffer containing 1.5% BSA and 0.05% Tween-20 in PBS for 1 hour at room temperature.

The embryos were stained overnight at 4°C with an anti-GATA4 antibody (sc- 1237, Santa Cruz, USA) diluted at 1:200. After being washed with blocking buffer three times, the secondary antibody, Alexa Fluor 488 donkey anti-goat IgG (1:500, A11055, Invitrogen), was used to stain the embryos for 1 hour in blocking buffer at room temperature. Nuclei were stained with 1 μg/ml Hoechst 33342 (19172-51, nacalai tesque, Japan) for 10 minutes. Then, embryos were washed twice in blocking buffer and mounted in 50% glycerol under micro cover glass.

